# CpG methylation accounts for genome-wide C>T mutation variation and cancer driver formation across cancer types

**DOI:** 10.1101/106872

**Authors:** Rebecca C. Poulos, Jake Olivier, Jason W. H. Wong

## Abstract

Cytosine methylation (5mC) is vital for cellular function, and yet 5mC sites are also commonly mutated in the genome. In this study, we analyse the genomes of over 900 cancer samples, together with tissue type-specific methylation and replication timing data. We describe a strong mutation-methylation association in colorectal cancers with microsatellite instability (MSI) or with *Polymerase epsilon (POLE)* exonuclease domain mutation. We describe a potential role for mismatch repair in the correction of mismatches resulting from deamination of 5mC, and propose a mutator phenotype to exist in *POLE*-mutant cancers specifically at 5mC sites. We also associate *POLE*-mutant hotspot coding mutations in *APC* and *TP53* with CpG methylation. Analysing mutations across additional cancer types, we identify nucleotide excision repair- and AID/APOBEC-induced processes to underlie differential mutation-methylation associations in certain cancer subtypes. This study reveals differential associations vital for accurately mapping regional variation in mutation density and pinpointing driver mutations in cancer.

## Introduction

Cancer develops as somatic mutations accumulate in cells, with certain driver mutations conferring a growth advantage to a sub-population (Nowell, 1976). In some cancers, mutations develop primarily from exposure to exogenous mutagens such as ultraviolet (UV) light or cigarette smoke, while in other cancer types, most mutations accumulate after a cell develops defective replication or repair mechanisms (Vogelstein et al., 2013). Mutation rates vary throughout the cancer genome due to factors such as trinucleotide composition (Alexandrov et al., 2013), transcription factor binding (Perera et al., 2016; Sabarinathan et al., 2016), replication timing, chromatin organisation (Schuster-Bockler and Lehner, 2012) and mismatch repair (MMR) efficiency (Supek and Lehner, 2015). However, the origin of many mutations within cancer cells still remains unknown (Alexandrov et al., 2013).

DNA methylation is an epigenetic mark most commonly occurring in the genome at sites of CpG dinucleotides (Riggs and Jones, 1983). Methylation involves the covalent attachment of a methyl group to the fifth atom of the carbon ring of a cytosine, forming molecules known as 5-methylcytosine (5mC) (Brero et al., 2006). Methylation has important functions within a cell, influencing development (Smith and Meissner, 2013), gene expression and silencing (Doerfler, 2006), as well as being implicated in carcinogenesis (Jones and Baylin, 2002).

Despite its crucial role in cellular function however, CpG methylation can also be somewhat mutagenic, with methylated cytosines being approximately fivefold more likely to undergo spontaneous deamination (loss of an amine group) than unmethylated cytosines (Ehrlich et al., 1986). 5mC deamination yields thymine, leading to a G•T mismatch in DNA which can be recognised by thymine DNA glycosylases and repaired through the base excision repair (BER) pathway (Jacobs and Schär, 2012; Walsh and Xu, 2006). However, if a cell replicates before the mismatch can be repaired, a C>T mutation will become encoded into its genome. A mutation signature from cytosine deamination at CpG sites (signatures 1A and 1B from Alexandrov et al. (2013)) has been identified in many cancer types, and is strongly correlated with age of diagnosis, as age allows more time for deamination events to occur and their effects to accumulate (Alexandrov et al., 2015). Methylated CpG dinucleotides (mCpGs) have additionally been found to be more highly mutated in non-cancer tissues, with mutation rates also correlating with increasing age (Rahbari et al., 2016).

The commonly accepted dogma regarding mutations at mCpGs is that mutations accumulate solely due to random spontaneous deamination of 5mC. However, other processes have also been associated with 5mC mutation or deamination, including exposure to UV light or to cigarette smoke (Pfeifer, 2006). In addition, understanding the repair of G•T mismatches is crucial in determining how mutations at sites of 5mC accumulate within the genome (Wiebauer et al., 1993). In this study, we analyse the association between methylation and mutation in 63 whole-genome sequenced (WGS) colorectal cancers, together with an additional 847 whole-genomes across 11 cancer types. We describe the association in detail within colorectal cancer subtypes, positing a potential role for MMR in the repair of deaminated 5mCs, and implicate *Polymerase epsilon* (*POLE*) exonuclease domain mutation (*POLE*-mutant) in increased mutagenesis at 5mC sites. We further define the influence of methylation and replication timing in driving the differential patterns of mutation accumulation and repair across the genomes of the additional cancer types and subtypes analysed, revealing associations of methylated CpG mutagenesis with nucleotide excision repair (NER) and deaminase enzyme activity.

## Results and Discussion

### Methylation and mutation associations in colorectal cancer

Recent studies investigating the accumulation of somatic mutations in cancer have shown that mutations in many cancer types increase at promoters due to inhibition of nucleotide excision repair (NER) at transcription factor bindings sites (Perera et al., 2016; Sabarinathan et al., 2016). Colon cancers were found to exhibit the lowest relative rate of mutations at promoters, attributable to the reduced importance of NER in the repair of mutations accumulating in such tissue (Perera et al., 2016). In this study, we investigate the reduction of promoter mutations in colorectal cancer further. To do so, we first constructed mutation profiles around transcription start sites (TSSs) using 63 WGS colorectal cancer samples from The Cancer Genome Atlas (TCGA), observing a decrease in mutation load in the region immediately surrounding the TSS (**Fig 1a**). To understand the effect across colorectal cancer subtypes, we separated our samples into those which were microsatellite stable (MSS), had microsatellite instability (MSI) or were *POLE*-mutant, finding each of the subtypes to exhibit reduced mutation loads at the TSS, with more pronounced relative hypo-mutation in MSI and *POLE*-mutant samples (**Supp Fig 1a**).

**Figure 1.**
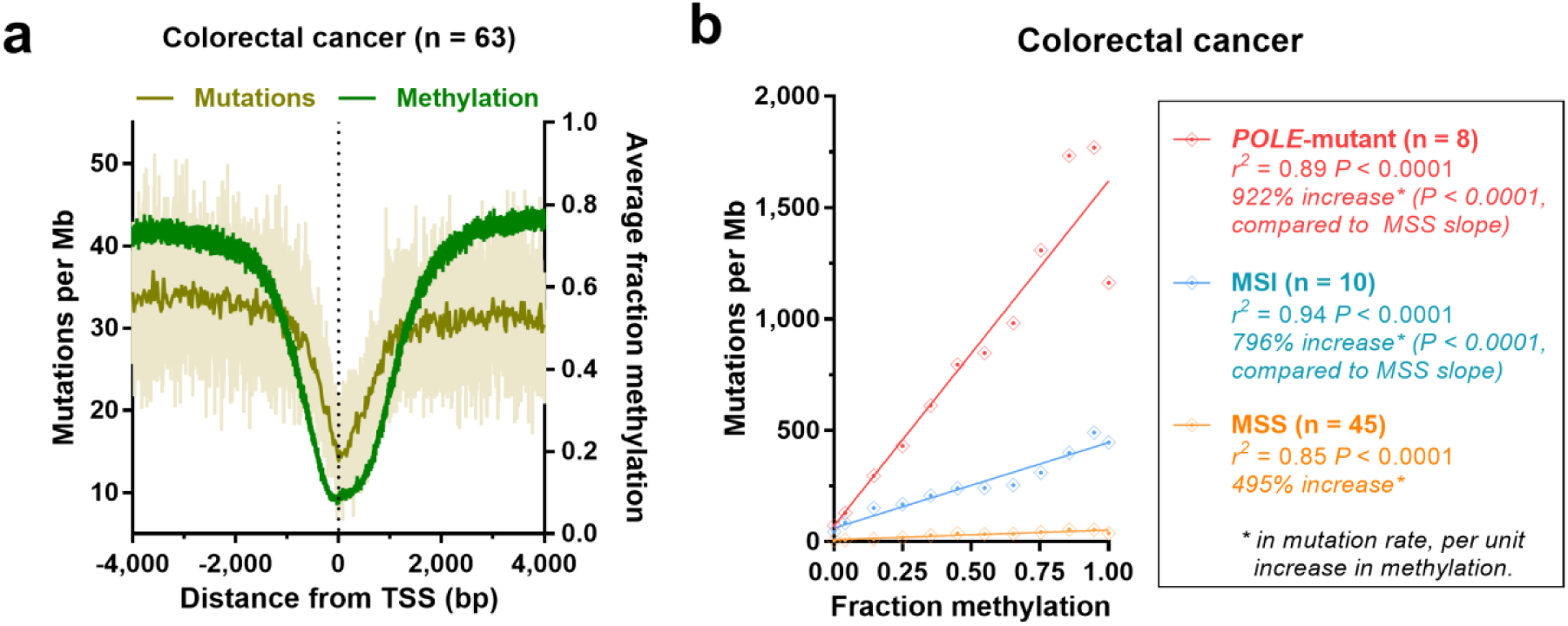
**Association between mutation accumulation and methylation in colorectal cancer subtypes. (a)** Colorectal cancer (n = 63) mutation profile and average methylation profile from colon whole genome bisulfite sequencing (WGBS) around transcription start sites (TSSs). Nucleotide-resolution mutation data (light beige), together with mutation data in 25 bp bins (dark beige) is shown. **(b)** Correlation between mutations and colon WGBS methylation across autosomes for *POLE*-mutant, MSI and MSS colorectal cancers. Binned data is shown (bins of 0.1 methylation), along with *r*^*2*^ and significance from Pearson’s regression. Mutation rate increase was calculated by Poisson regression on binned data, with MSS as the reference factor.

As CpG methylation is typically lower at CGI-associated promoter elements (Long et al., 2016), we associated methylation and mutations around the TSS, using normal sigmoid colon whole-genome bisulfite sequencing (WGBS) data (Roadmap Epigenomics Consortium et al., 2015). We mapped average CpG methylation, observing a corresponding decrease in methylation in the region immediately surrounding the TSS (**Fig 1a**; see also **Supp Fig 1b** for DNase I hypersensitivity (DHS) and H3K4me3 profiles around the TSS – indicating promoter activity). We measured the association between methylation and CpG mutations per Mb across autosomes in colorectal cancer, finding a significant association for each subtype (*P* < 0.0001, Pearson’s correlation; **Fig 1b**). We observed significantly greater slopes in *POLE*-mutant and MSI samples when compared with the slope in MSS samples (72% and 51% greater rate of mutation per unit increase in methylation respectively, *P* < 0.0001, Poisson regression; **Fig 1b**). This finding shows that despite the increased mutation load of MSI and *POLE*-mutant colorectal cancers, mutations at CpG sites remain methylation-associated. Taking the mutation-methylation association in MSS samples to be that occurring due to endogenous mCpG deamination and repair in colon tissue, the greater slopes of MSI and *POLE*-mutant cancers must be attributable either to increased mutagenesis or to deficiencies of repair.

### Potential role for mismatch repair in 5mC deamination repair and regional variation

At megabase scales, MMR activity causes genome-wide variation in mutation load according to replication timing (Schuster-Bockler and Lehner, 2012; Supek and Lehner, 2015). This means that in MSI colorectal cancers (which are MMR-deficient), genomic variation at such scales is mostly lost (Supek and Lehner, 2015). We note that average methylation, varies only slightly with replication timing changes at megabase scales (*r*^*2*^ = 0.20 P < 0.0001, Pearson’s correlation;  = 0.0013, with fraction methylation per line of best fit, ranging from 0.75 to 0.83; **Supp Fig 2a**). We hypothesised therefore, that the strong mutation-methylation association in MSI colorectal cancers (see **Fig 1b**) may underlie the reported loss of genomic variability (Supek and Lehner, 2015), with MMR during replication playing a significant role in the repair of the G•T mismatches that result from 5mC deamination.

To investigate this hypothesis, we isolated C>T mutations at CpGs in MSS and *POLE*-mutant colorectal cancers. We also isolated C>T CpG mutations from late-onset MSI (median 66.7% of time spent as MSI) and early-onset MSI (median 84.3% of time spent as MSI) samples (*P* < 0.0001, unpaired t-test; **Supp Fig 2b**), boosting sample sizes by incorporating WGS uterine and stomach cancers into our MSI subsets. We then correlated mutations with replication timing, finding a positive correlation in both MSS and *POLE*-mutant colorectal cancers (*r*^*2*^ = 0.97 *P* < 0.001 and *r*^*2*^ = 0.64 *P* = 0.0567 respectively, Pearson’s correlation; **Fig 2a**), consistent with the higher rates of mutation accumulation known to occur in late-replicating regions (Schuster-Bockler and Lehner, 2012; Supek and Lehner, 2015). Strikingly however, we found both early- and late-onset MSI cancers to exhibit significant negative correlations between C>T mutations and replication timing at CpG dinucleotides (*r*^*2*^ = 0.92 and *r*^*2*^ = 0.88 respectively, *P* < 0.01, Pearson’s correlation; **Fig 2a**), with more mutations occurring at early-replicating regions. In fact, the earlier that a sample gained MSI, the more negative was the slope of the line of best fit between CpG C>T mutations and replication timing (*r*^*2*^ = 0.50 *P* < 0.05, Pearson’s correlation; **Supp Fig 2c**).

**Figure 2.**
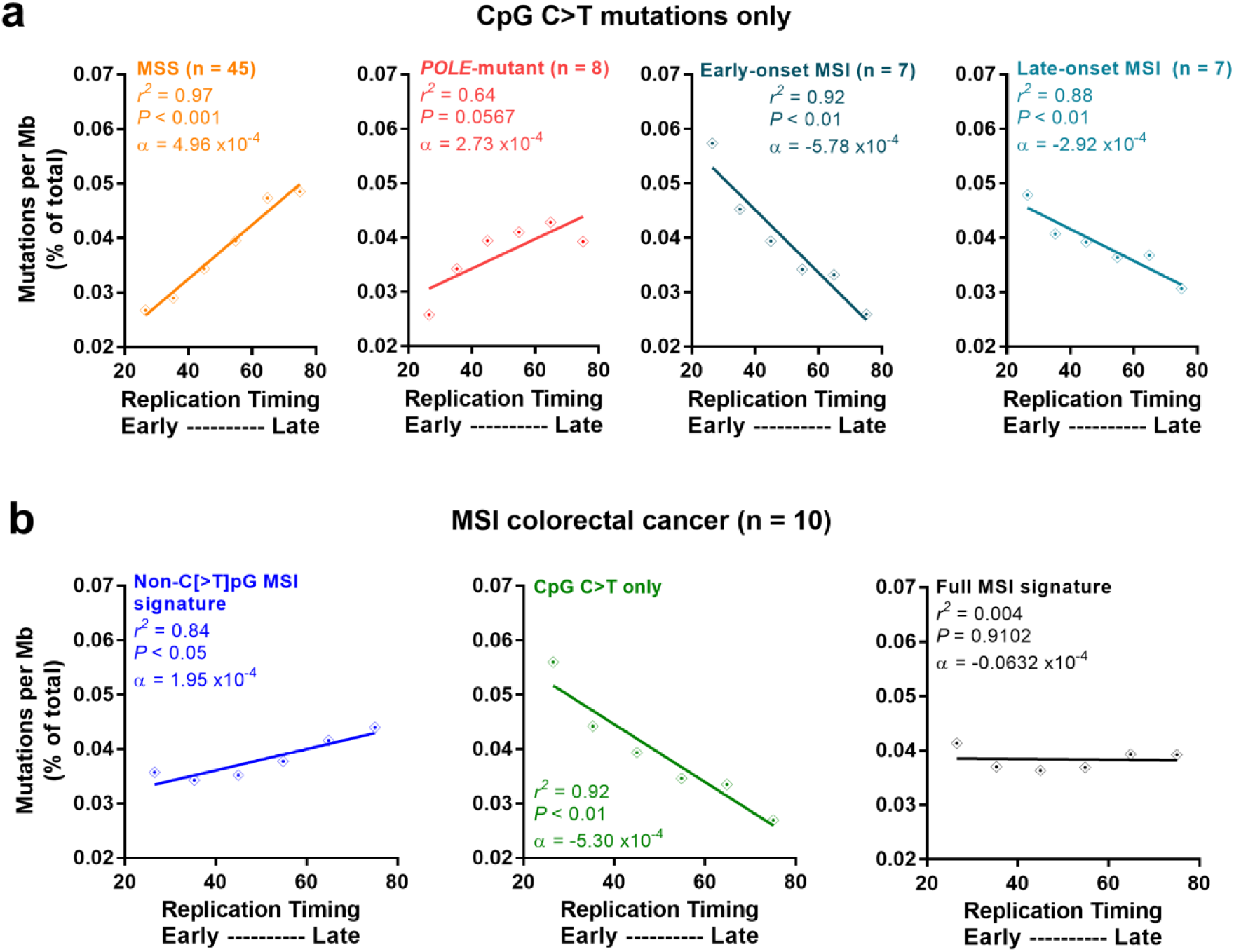
**Replication timing and MSI-associated mutations in colorectal cancer subtypes (a)** Association between CpG C>T mutations and replication timing for microsatellite stable (MSS) colorectal cancers, and those with *Polymerase epsilon* exonuclease domain mutation (*POLE*-mutant). Association also shown for colorectal, uterine and stomach cancers with early– and late-onset of microsatellite instability (MSI). **(b)** Association between non-C[>T]pG MSI signature, CpG C>T, and full MSI signature (including CpG C>T) mutations with replication timing for MSI colorectal cancers. Data points show binned data (bins of 10 replication timing), with *r*^*2*^ and significance by Pearson’s correlation. *α* denotes the slope of the line.

*In vivo* studies using SV40 heteroduplex transfection have shown that MMR is involved in the repair of a small proportion (~8%) of 5mC deamination events (Brown and Jiricny, 1987, 1988; Wiebauer et al., 1993). Our findings also suggest that MMR may play a role in the repair of deamination-induced G•T mismatches. The increased time available for BER to recognise and repair the G•T mismatches resulting from 5mC deamination prior to cellular replication should result in a decrease in mutation load in late-replicating regions. In MSI samples, such an association is evident (**Fig 2a**), with most CpG mutations accumulating in the absence of MMR. In contrast, since MMR activity is enhanced in early-replicating euchromatic regions (Supek and Lehner, 2015), when MMR is proficient – as in MSS and *POLE*-mutant colorectal cancers – we see a decrease in mutation load in such regions (**Fig 2a**), obscuring what may be an otherwise negative BER-associated trend. We find no evidence for decreased BER function in MSI cancers (see **Supplementary Data**), leading us to suggest MMR involvement in the repair of mismatches at mCpG sites, with the precise mechanism requiring further research.

### mCpG mutability contributes to the regional loss of mutation density variation in MSI cancers

To determine whether the negative correlation that we observed in **Fig 2a** does contribute to the reported overall genome-wide loss of mutation density variation in MSI samples (Supek and Lehner, 2015) as we had hypothesised, we plotted the correlation of non-C[>T]pG MSI signature mutations (see **Methods**), CpG C>T mutations and full (CpG-inclusive) MSI signature mutations against replication timing (**Fig 2b**). The full MSI signature (*n* = 323,848 mutations in MSI samples) is thus the combination of both non-C[>T]pG MSI signature (*n* = 236,732; 73%) and CpG C>T (*n* = 87,116; 27%) mutations. Our analysis confirmed the findings of Supek & Lehner (2015), as the slope of the mutation-replication timing association was consistently lower across all mutation categories in MSI samples when compared with MSS and *POLE*-mutant samples (**Fig 2b**, **Supp Fig 2d**). However, our analyses also showed that the negative association of CpG C>T mutations in MSI cancers (*α* = −5.30x10^-4^) does contribute to the magnitude of genomic variability loss according to replication timing for overall CpG-inclusive MSI-associated mutations (*α* = −0.0632x10^-4^; non-C[>T]pG MSI signature mutations *α* = 1.95x10^-4^; **Fig 2b**). Thus, while MMR does underlie mutation rate variation across the genome, we suggest that the effect of MMR deficiency on mutation accumulation at CpG dinucleotides is a factor that contributes to the *extent* of this loss of mutation-replication timing-association.

### Mutagenesis at 5mC nucleotides in POLE-mutant colorectal cancers

We computed the correlation coefficient between CpG mutations and methylation separately for individual *POLE*-mutant colorectal cancer samples, finding the slope of the line of best fit from binned data to range from 19.23 to 3,090 (**Supp Fig 3**). We found the slopes to significantly positively correlate with the total number of mutations in each *POLE*-mutant sample (*r*^*2*^ = 0.75 *P* < 0.01, Pearson’s correlation; **Fig 3a**), suggesting that much of the increased mutagenesis at CpGs in *POLE*-mutant cancers is methylation-associated. *POLE*-mutant samples have an inactivated exonuclease domain, leading to a loss of proofreading ability on newly-synthesized DNA (Kane and Shcherbakova, 2014; Rayner et al., 2016). Samples with greater absolute numbers of mutations therefore likely have either a stronger mutator phenotype, or have become *POLE* exonuclease domain mutated earlier. However, with neither of these factors expected to alter the rate of 5mC deamination, we hypothesised that exonuclease domain-mutated POLE may more often make replication errors when encountering a site requiring the insertion of guanine in a mCpG context. It is possible that these data could also be explained if errors are introduced by wild-type POLE when encountering a mCpG context and, where there is proofreading deficiency, these errors remain uncorrected in the genome. However, to our knowledge, there is no evidence from *in vitro* studies that wild-type replicative polymerases typically make such errors in the context of methylated cytosines.

**Figure 3.**
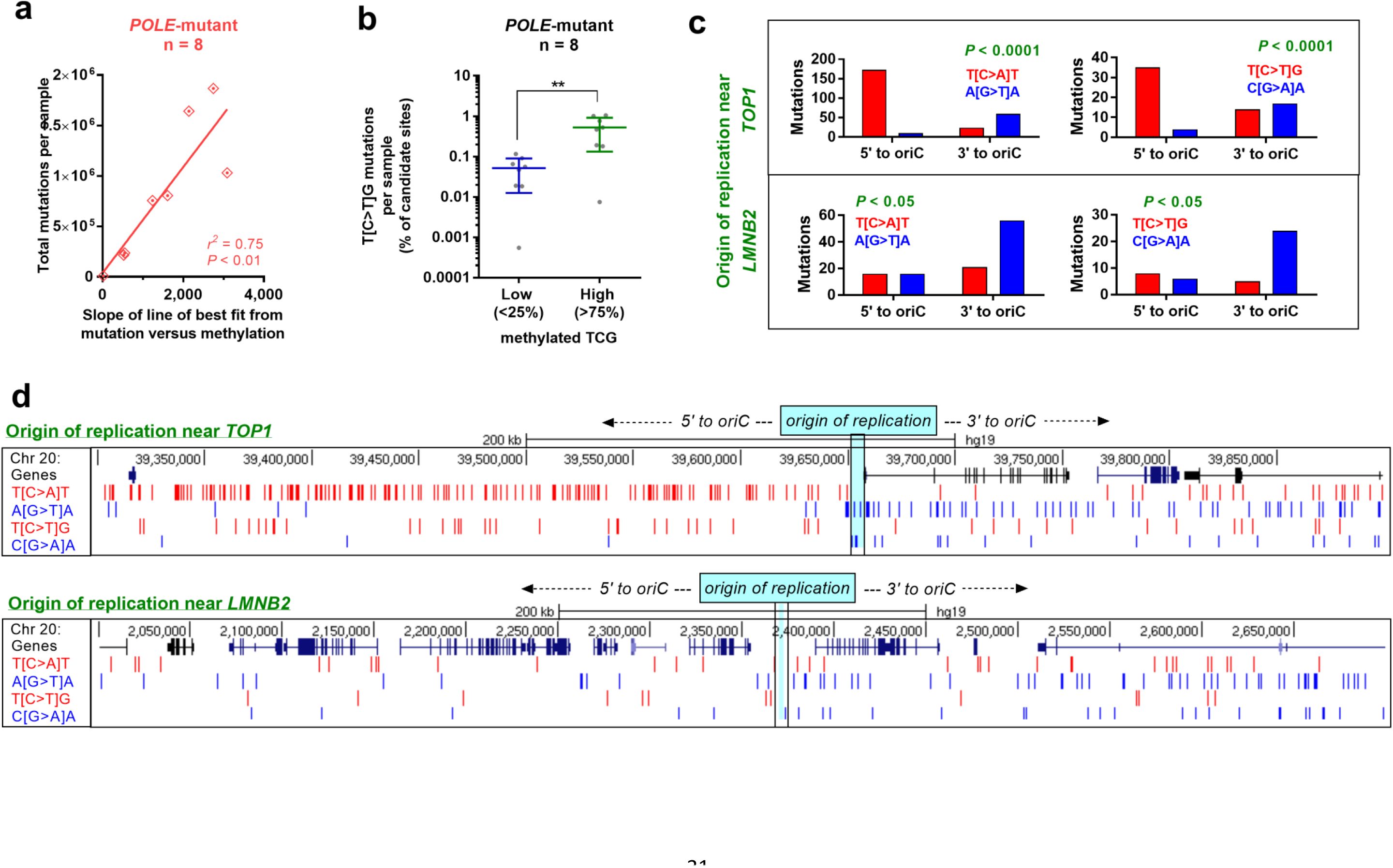
**Methylation-associated mutations in *POLE*-mutant colorectal cancers. (a)** Correlation of total mutations per *Polymerase epsilon* exonuclease domain mutant (*POLE*-mutant) colorectal cancer sample, with the slope of line of best fit from the mutation-methylation association at Supp Fig 3. *r*^*2*^ and significance is by Pearson’s correlation. (b) Percentage of candidate sites which harbour C>T mutations in a TCG context per sample for low (<25%) and high (>75%) methylated CpGs (in normal colon tissue) per *POLE*-mutant colorectal cancer sample. Significance is by unpaired t-test where ** *P* < 0.01. (c) Strand-specificity of T[C>A]T (left) and T[C>T]G (right) mutations in the regions 5’ and 3’ to origins of replication (oriC) near *TOP1* (top) and *LMNB2* (bottom). Significance is by Fisher’s exact test. (d) Excerpt from the UCSC genome browser, depicting strand specificity of T[C>A]T and T[C>T]G mutations 5’ and 3’ to the oriC near *TOP1* (top) and *LMNB2* (bottom).

Consistent with our hypothesis of a *POLE*-mutant mutator phenotype, we found a significantly greater proportion of T[C>T]G mutations to occur at high rather than low methylated TCG sites (*P* < 0.01, paired t-test; **Fig 3b**), with the TCG trinucleotide being the most highly mutated CpG variant in *POLE*-mutant tumours (Alexandrov et al., 2013; Shinbrot et al., 2014). Given POLE’s role in leading strand replication (Miyabe et al., 2011; Pursell et al., 2007), we investigated the strand-specificity of the T[C>A]T and T[C>T]G mutations, which are both common mutations in *POLE*-mutant cancer genomes (Alexandrov et al., 2013; Shinbrot et al., 2014). In support of our mutator-phenotype hypothesis, we found significant strand asymmetry to occur in both trinucleotide contexts around known origins of replication (*P* < 0.05 and *P* < 0.0001, Fisher’s exact test; **Fig 3c**, **d**). There is a growing body of evidence which suggests that *POLE* exonuclease domain mutation can result in a mutator phenotype greater than that from proofreading-deficiency alone (Kane and Shcherbakova, 2014). Some variants have been shown to increase mutation load above that from inactivation of the catalytic domain alone (Shinbrot et al., 2014), lending further support to our hypothesis.

### mCpG mutations as potential driver events in POLE-mutant colorectal cancers

Many mutations responsible for genetic diseases are C>T transitions occurring at CpG dinucleotides (Cooper et al., 2011; Walsh and Xu, 2006). Additionally, methylated CpGs are hotspots for somatic cancer mutations in driver genes such as *TP53, RB1* and *EGFR* (Fujii et al., 2015; Holliday and Grigg, 1993; Jones et al., 1992; Walsh and Xu, 2006). *POLE*-mutant colorectal cancers harbour specific mutation hotspots in the key tumour-suppressors *TP53* and *APC* (Palles et al., 2013; Shinbrot et al., 2014) (a finding which we have confirmed in our samples (**Fig 4a**)). As *POLE* exonuclease domain mutation is thought to be an early event in tumours (Rayner et al., 2016), these *POLE*-mutant-signature mutations could occur reasonably early in oncogenesis, serving as gatekeeper mutations which confer a growth advantage to cellular subpopulations, and driving tumour growth. We observed that these mutation hotspot (truncating C>T mutations at *TP53* R213X and *APC* R1114X) occur at TCG trinucleotides, and so we hypothesised that these sites may be more often mutated specifically in *POLE*-mutant tumours because of the strong mutation-methylation association in this subtype (see **Fig 1b**).

**Figure 4.**
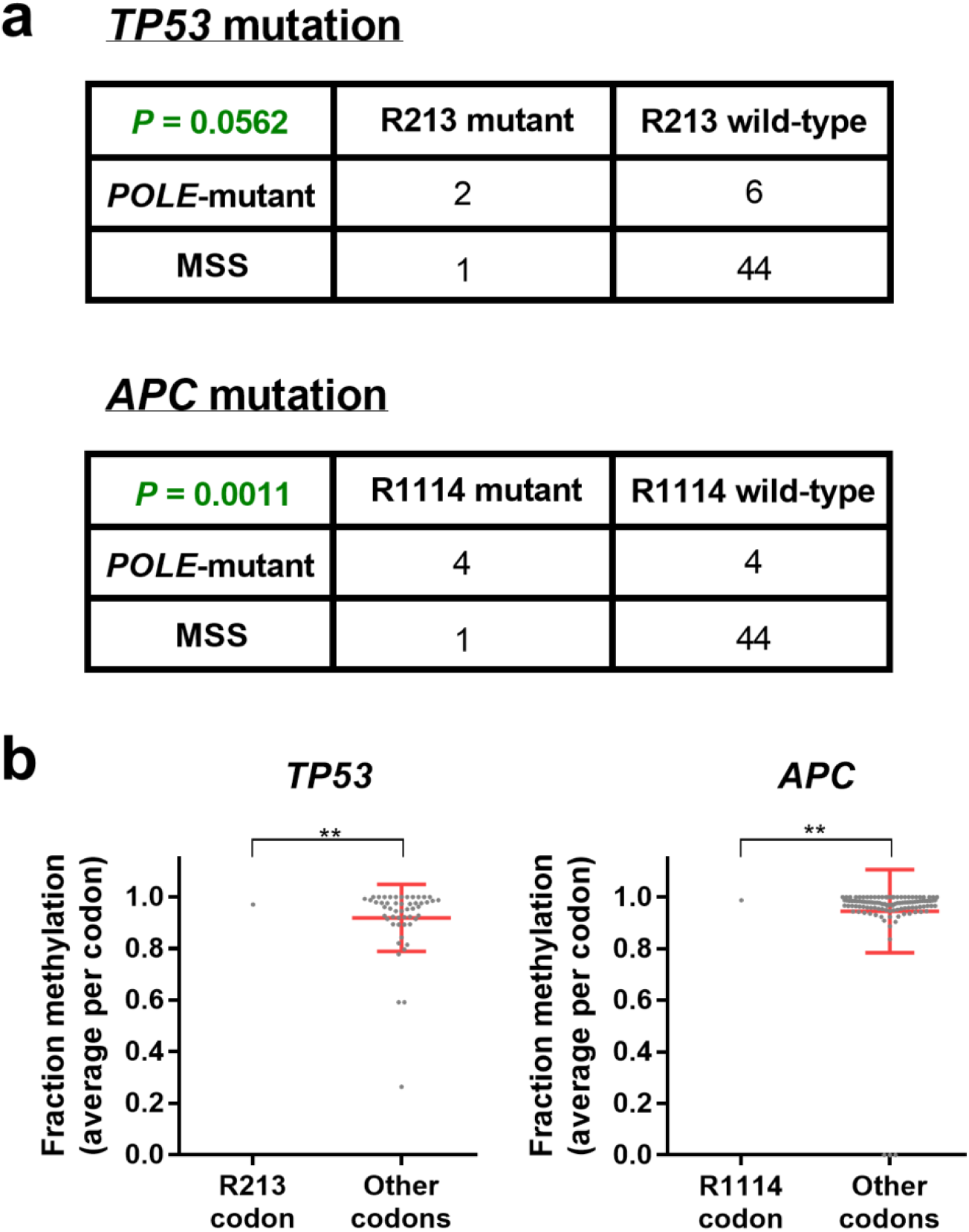
**Mutation hotspots in cancer driver genes in *POLE*-mutant tumours. (a)** Contingency table and significance from Fisher’s exact test of *Polymerase epsilon* exonuclease domain mutant (*POLE*-mutant) and microsatellite stable (MSS) colorectal cancer samples which are wild-type or mutant at *TP53* R213 and *APC* R114 codons. (b) Methylation status in normal colon tissue for each CpG site within coding exons of *TP53* and *APC*, together with significance by one-sample t-test against methylation at R213 and R1114 codons respectively. ** denotes *P* < 0.01.

We found that these sites are in fact highly methylated in normal colon tissue, with the CpG at *TP53* R213 methylated in 97.1% of cells, and at *APC* R1114 methylated in 98.7% of cells (**Fig 4b**). However, while these sites are methylated to a significantly greater extent than other codons in the same gene (*P* < 0.01, one-sample t-test; **Fig 4b**), there may yet be other locations in *TP53* or *APC* which are equally likely to become mutated – when considering methylation alone. To investigate this, we considered all possible C>T mutations at TCG trinucleotides which would lead to the immediate truncation of either *TP53* or *APC*. We found that the R213 site in *TP53* is the only possible trinucleotide which fulfils these criteria (**Supp Fig 4a**), adequately explaining its hotspot mutation status in *POLE*-mutant samples. In *APC* however, we found three additional sites occurring earlier from the N-terminal of the protein which fulfilled the criteria listed, together with five mutation sites at or after the C-terminal of codon 1920 (**Supp Fig 4b**). This suggests that while methylation may be responsible for the formation of specific mutation hotspots in *POLE*-mutant cancers, other factors may also contribute to mutation occurrence and selection within cells – perhaps due to a phenotype conferred to cells by mutations at specific sites, making them more likely to be observed in cancer sequencing data (Walsh and Xu, 2006).

### Differential influence of methylation on mutation accumulation across cancer types and subtypes

Having described a strong mutation-methylation association across colorectal cancer subtypes, we next sought to investigate whether any such association exists in other cancer types. To do so, we incorporated into our analyses, somatic mutations from an additional 847 cancer samples across 11 cancer types available from TCGA and previously published datasets (Zheng et al., 2014). We developed regression models using both tissue type-specific methylation data (**Supp Table 1**) and average cell-type replication timing data, plotting actual mutations together with the function predicted by multivariable regression models (see **Methods**).

To first validate our regression models, we investigated the predicted associations in colorectal cancer, finding a positive association between mutation probability and fraction methylation across colorectal cancer subtypes for all possible methylation values (function vertex > 1; **Table 1**), consistent with what we have already demonstrated (see **Fig 1b**). Also confirming previous findings (Supek and Lehner, 2015), we found mutation probability to vary little across replication timing changes in MSI colorectal cancers, compared with MSS and *POLE*-mutant subtypes (depicted in rightmost graphs, **Fig 5a-c**). This is demonstrated by the small improvement to the area under the curve (AUC) in nested models which additionally incorporated methylation (MSI: 2.0%), compared with 16.5% in MSS and 12.5% in *POLE*-mutant subtypes (**Table 1**).

**Table 1.** Regression equation from multivariable models predicting mutation probability across colorectal and skin cancer subtypes, together with vertex and area under curve (AUC) predictions.

| Cancer type/subtype | Regression model equation* | Vertex<sup>^</sup> (fraction methylation) | AUC<sup>@</sup> for regression model A<sup>A</sup> | AUC<sup>+</sup> for regression model B<sup>B</sup> | AUC<sup>+</sup> for regression model C<sup>C</sup> | Increase to AUC via replication<sup>#</sup> | Increase to AUC via methylation<sup>+</sup> |
| --- | --- | --- | --- | --- | --- | --- | --- |
| MSS colorectal cancer | $y = -5.788 + 3.2436M + -1.3425M^2 + -0.0277R + -0.0026(M \times R)$ | 1.21 | 0.579 | 0.657 | 0.674 | 16.5% | 2.7% |
| MSI colorectal cancer | $y = -5.7974 + 1.6834M + -0.1411M^2 + -0.0142R + 0.0082(M \times R)$ | 5.97 | 0.589 | 0.540 | 0.600 | 2.0% | 11.1% |
| *POLE*-mutant colorectal cancer | $y = -5.0593 + 4.7117M + -2.2413M^2 + -0.0265R + 0.0058(M \times R)$ | 1.05 | 0.580 | 0.619 | 0.653 | 12.5% | 5.5% |
| Melanoma | $y = -3.5202 + 3.6233M + -3.5961M^2 + -0.0379R + 0.0117(M \times R)$ | 0.50 | 0.595 | 0.659 | 0.674 | 13.4% | 2.4% |
| XPC<sup>-/-</sup> squamous cell carcinoma | $y = -5.6325 + 4.8751M + -3.7947M^2 + -0.0232R + 0.008(M \times R)$ | 0.64 | 0.586 | 0.600 | 0.630 | 7.4% | 4.9% |

\* See **Methods** for canonical regression model formula, where y = log odds of mutation probability, M = methylation and R = replication timing
<sup>^</sup> Vertex (unit: fraction methylation) predicted by regression model, calculated as $-b_1/(2 \times b_2)$ .
<sup>@</sup> AUC = area under curve
<sup>A</sup> Regression model equation: $y = b_0 + b_1 M + b_2 M^2$ where y = log odds of mutation probability, M = methylation and R = replication timing
<sup>B</sup> Regression model equation: $y = b_0 + b_1 R$ where y = log odds of mutation probability, M = methylation and R = replication timing
<sup>C</sup> Regression model equation: $y = b_0 + b_1 M + b_2 M^2 + b_3 R$ , where y = log odds of mutation probability, M = methylation and R = replication timing
<sup>#</sup> Calculated using values of AUC from models: (C-A)/A
<sup>+</sup> Calculated using values of AUC from models: (C-B)/B

**Figure 5.**
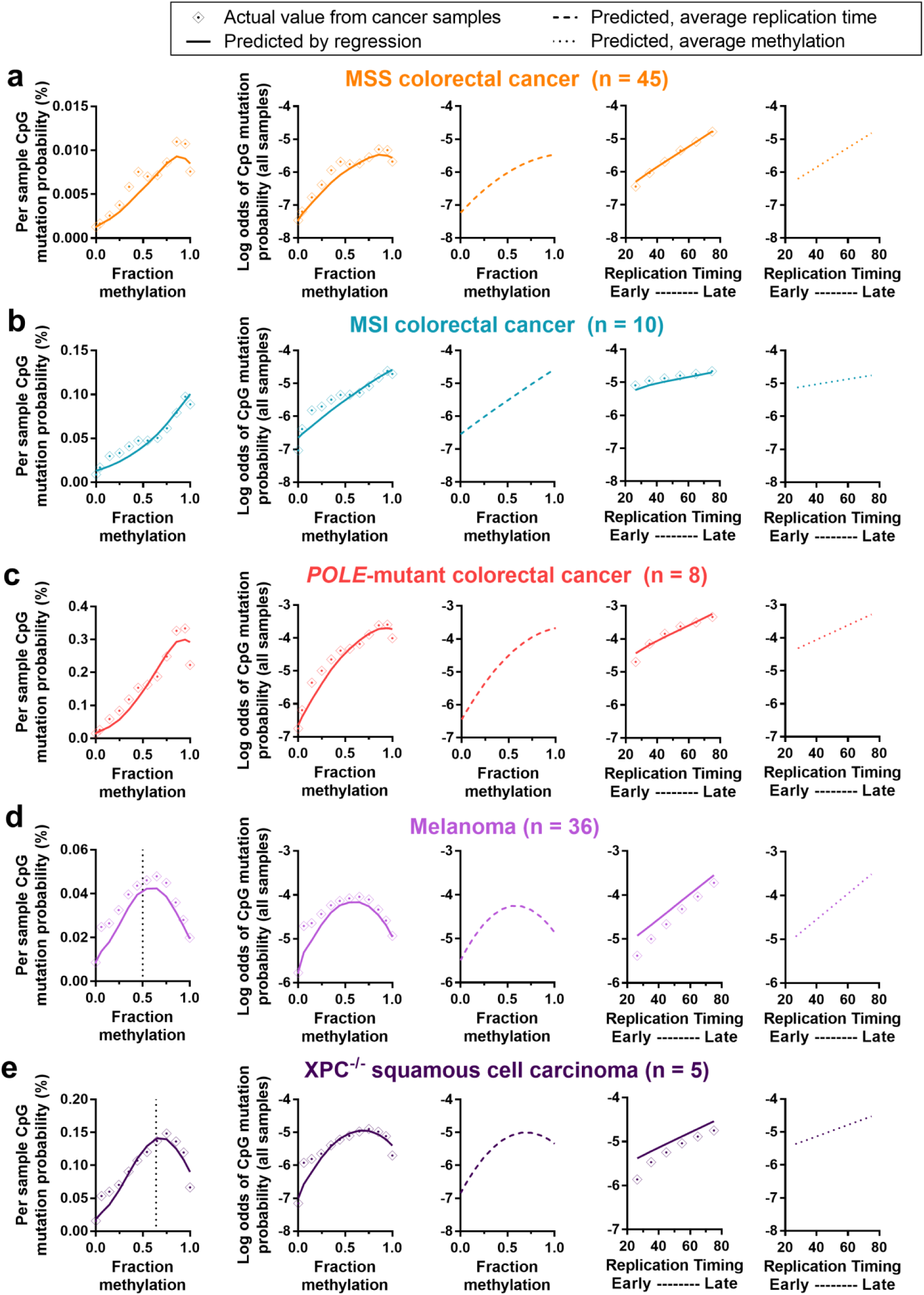
**Actual and predicted mutation rates, according to methylation and replication timing, for colorectal and skin cancer subtypes.** Graphs depict actual and predicted (by regression model; see Methods) mutation probability and log odds of mutation probability by methylation or replication timing, for (a) microsatellite stable (MSS) colorectal cancer, (b) colorectal cancers with microsatellite instability (MSI), (c) colorectal cancers with *Polymerase epsilon* exonuclease domain mutation (*POLE*-mutant), (d) melanoma and (e)XPC^-/-^ squamous cell carcinoma. Graphs from left to right are: mutation probability by fraction methylation (actual and predicted), log odds of mutation probability by fraction methylation (actual and predicted), log odds of mutation probability by fraction methylation (predicted, using overall average replication timing in all bins), log odds of mutation probability by replication timing (actual and predicted) and log odds of mutation probability by replication timing (predicted, using overall average methylation in all bins). Binned data is shown (bins of 0.1 for methylation or 10 for replication timing), with any vertex between 0 and 1 fraction methylation indicated by a dotted line. See Table 1 for regression output, predicted vertex and area under curve values.

We next examined the mutation-methylation association in skin cancer subtypes, as skin cancers are subject to well-defined mutation and repair processes associated with UV light. Interestingly, we found that the association between mutation rate and methylation was not positive across all methylation values (**Fig 5d, e**). In melanoma, the vertex predicted by the multivariable regression model was at 0.50 fraction methylation (**Table 1**), meaning that at methylation fractions greater than 0.50, increasing methylation was associated with decreasing mutation probability (leftmost graph; **Fig 5d**). When removing replication timing variation from the function (see **Methods**), the association between methylation and the log odds of mutation probability remained negative at high levels of methylation (middle graph; **Fig 5d**) possibly suggesting either that this is the true underlying association, or that currently unknown factors are influencing the mutation rate at highly methylated CpGs in melanoma.

NER is particularly vital in skin cancers due to its role in the repair of UV light-induced DNA lesions (Schärer, 2013). We found replication timing and levels of NER in response to UV light exposure (Zheng et al., 2014) to be significantly correlated with one another (cyclobutane pyrimidine dimer (CPD): *r*^*2*^ = 0.79, and (6-4)pyrimidine-pyrimidone photoproduct ((6-4)PP): *r*^*2*^ = 0.59, *P* < 0.0001, Pearson’s correlation; **Supp Fig 6a**). Strikingly, we also found that the mutation-methylation pattern in melanoma closely mimics the replication timing-methylation pattern in NHEK cells (**Supp Fig 6b**), likely attributable to the association of replication timing and chromatin density. The propensity for mutagenic CPD DNA lesion formation following UV light exposure is known to increase at mCpGs (Cannistraro et al., 2015; Rochette et al., 2009), and hence we expect that there is an underlying positive linear association between CpG mutation rate and methylation in UV light-induced cancers. If this is true, then this linear association should become clearer in XPC^-/-^ squamous cell carcinomas (SCCs), as XPC^-/-^ SCCs are global genome NER-deficient (Zheng et al., 2014) and ought to have a mutation profile mostly absent the influence of any NER-induced replication timing-association. Confirming this expectation, we found the vertex of the function predicting mutation probability to be at 0.64 fraction methylation in XPC^-/-^ SCC (**Table 1**), meaning that the mutation-methylation association remained positive over a greater range of methylation values in XPC^-/-^ SCC than it did in melanoma. Further, the AUC showed a 4.9% improvement when methylation was added to a nested model, with only a 2.4% improvement in melanoma (**Table 1**). Some highly methylated regions are active gene bodies which tend to be both early-replicating (Aran et al., 2010) and subject to transcription-coupled NER (Zheng et al., 2014), possibly leading to their reduced overall mutation load in melanoma. Taken together, our results suggest that the negative association between mutation rate and methylation at high fractions of methylation may, at least in part, be driven by the underlying mutation-replication timing-association.

When investigating other cancer types, we found that multivariable regression models predicted the function’s vertex to be between 0 and 1 fraction methylation for some cancers. This was the case in breast, liver, ovarian and pancreatic cancers, as well as chronic lymphocytic leukaemia (**Supp Table 2**). For these cancers, like in skin cancer, the association between mutation probability and fraction methylation was negative at some higher values of methylation (see **Supp Fig 5**). The primary mutation and repair processes are not well understood in many of these cancers, with samples harbouring varied mutation signatures and many mutations of unknown origin (Alexandrov et al., 2013). It is possible that our regression models are unable to completely separate the association between replication timing and methylation, with both factors significantly impacting on mutation rate. However, it may also be true that in some cancer types, the underlying association with methylation is actually such that, at high rates of methylation, mCpGs are less likely to become mutated. This may be due to the specific mutation and repair processes inherent in various tissue types, which are not well understood. In fact, other analyses have shown that the genome-wide rate of C>T single nucleotide polymorphisms (SNPs) increases only at low and intermediate (20-60%) methylation levels, but not at highly methylated sites (Xia et al., 2012).

### Influence of methylation on mutation accumulation in cancers with AID/APOBEC mutation signature

Many cancer types harbour mutation signatures related to the action of activation induced deaminases (AID) or apolipoprotein B mRNA editing enzyme catalytic polypeptide-like (APOBEC) enzymes (Alexandrov et al., 2013), a family of DNA deaminases involved in immunity and DNA demethylation (Rebhandl et al., 2015). These enzymes have been found to have substantially altered activity on differentially methylated cytosines (Nabel et al., 2012), and hence we investigated mutation rates at mCpG dinucleotides in breast cancer samples with and without mutation signatures indicating AID/APOBEC enzyme activity (see **Methods**). We found that regression models predicted the function’s vertex to shift from 0.75 fraction methylation in breast cancer samples without AID/APOBEC enzyme activity signatures to 0.50 fraction methylation in samples with those signatures (**Fig 6b**; **Supp Table 2**). This means that in breast cancer samples with AID/APOBEC enzyme activity signatures, mutation probability decreases as methylation increases, across a much greater range of methylation values than it does in samples without the signatures. This observation supports findings from biochemical studies indicating reduced activity of AID/APOBEC enzymes on 5mC (Nabel et al., 2012), translating such studies into an *in vivo* biological context.

**Figure 6.**
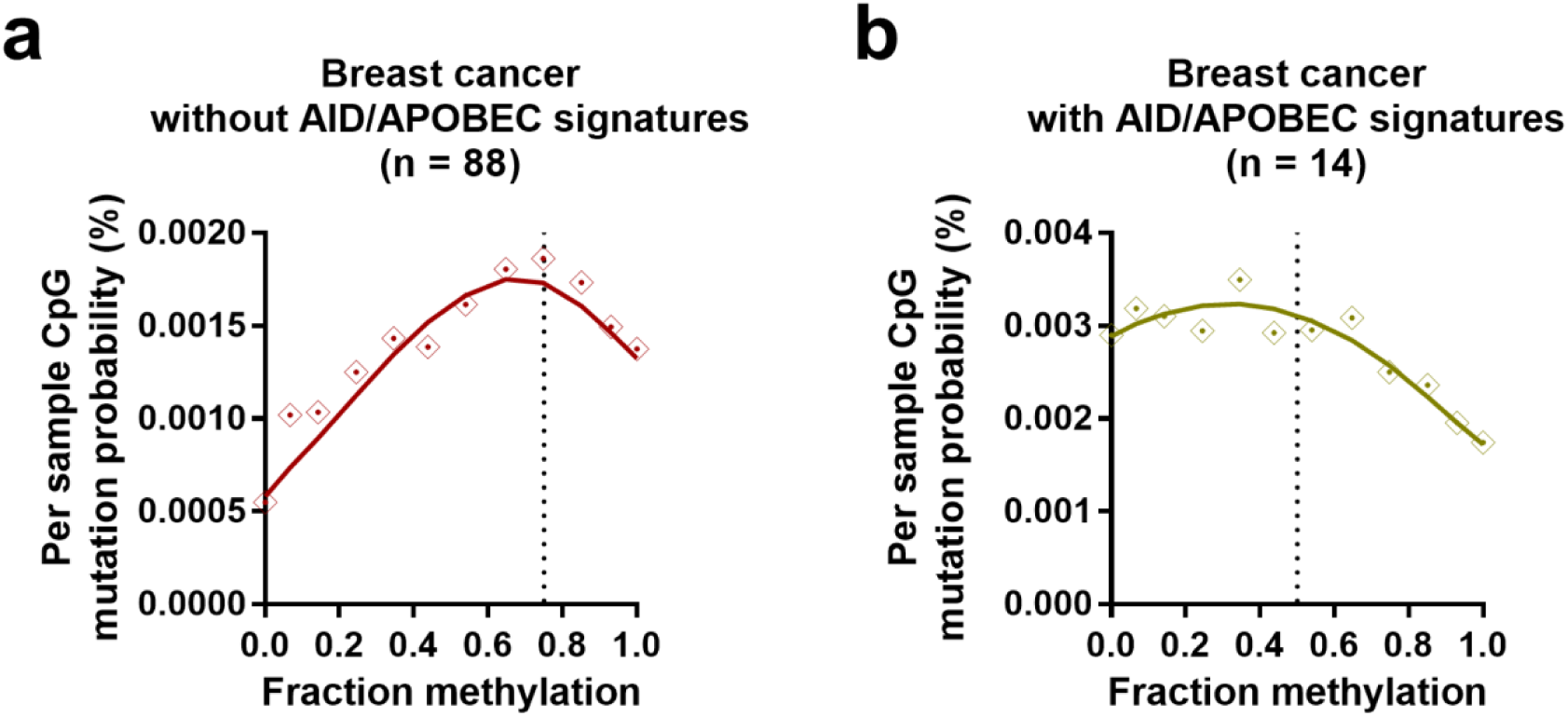
**Association between methylation and mutation probability in breast cancer samples with and without AID/APOBEC mutations signatures.** Graphs depict actual and predicted (by regression model; see Methods) mutation probability against fraction methylation for breast cancer samples (a) without and (b) with AID/APOBEC enzyme deaminase mutation signatures. Binned data is shown (bins of 0.1 for methylation) with any vertex between 0 and 1 fraction methylation indicated by a dotted line. See **Supp Table 2** for regression output, predicted vertex and area under curve values.

## Conclusion

In this study, we analyse 63 colorectal cancer whole-genomes, together with data from an additional 11 cancer types. Using tissue-specific methylation data, we describe a strong association between C>T mutations and methylation at CpG dinucleotides in many cancer types, driving patterns of mutation formation throughout the genome. Using analyses of replication timing, we suggest a potential role for MMR in the repair of G•T mismatches resulting from deamination of 5mC. We also propose a specific mutator phenotype to exist at methylated CpGs resulting from *POLE* exonuclease domain mutation – a phenotype which may be responsible for driving tumour growth through the formation of specific mutation hotspots in key cancer-associated genes. Additionally, we reveal differential associations between mutations and methylation across cancer types, with our findings providing significant developments in our understanding of mutation formation and repair at CpG dinucleotides in cells.

These analyses highlight the importance of understanding differential methylation-mutation associations across cancer types and subtypes. We emphasise the need for researchers to understand and stratify cancer subtypes according to their underlying mutation and repair processes when developing predictive models of expected mutation loads according to genetic and epigenetic features in the genome. Our work may assist cancer researchers in more accurately identifying driver mutations from among passenger mutations, by further defining expected background mutation rates in cancer genomes.

## Methods

### Somatic alterations and sample subtype classification

Somatic point mutations calls were made from BAM files available from The Cancer Genome Atlas (TCGA), as previously described (Perera et al., 2016), with the exception of mutation calls for XPC^-/-^ squamous cell carcinoma, which were taken directly from Zheng et al. (2014).

MSI samples were selected if they were listed as “MSI-H” in TCGA “clinical” data from the TCGA data portal. Early-and late-onset MSI samples were obtained from a previous study (Supek and Lehner, 2015), with additional WGS ‘UCEC’ and ‘STAD’ samples from TCGA. The early-onset MSI samples used in analyses were colorectal cancer samples TCGA-A6-6781, TCGA-AD-6964 and TCGA-AA-A00R, together with UCEC samples TCGA-AP-A0LD, TCGA-B5-A11H and TCGA-AP-A054 and STAD sample TCGA-CG-5723. The late-onset MSI samples used in analyses were colorectal cancer samples TCGA-QG-A5Z2, TCGA-D5-6540, TCGA-AA-3518 and TCGA-AA-3516, together with UCEC samples TCGA-A5-A0G9 and TCGA-BS-A0TE and STAD sample TCGA-BR-4280.

*POLE*-mutant samples were selected as such if they had a genome-wide mutation signature correlation with Signature 10 (Alexandrov et al., 2013) of greater than 0.85. Sample classifications were confirmed as exonuclease domain mutated if they contained an exonuclease domain mutation (between codons 268-471) as listed by Shinbrot et al. (2014) in “Supplemental Table 1A”.

To select breast cancer samples with and without AID/APOBEC signatures, deconstructSigs (Rosenthal et al., 2016) was run on each sample. Breast cancer samples were determined to have AID/APOBEC signatures if they had greater than 5% of mutations designated each to signatures 2 and 13, as these signatures have been attributed to the activity of AID/APOBEC family of cytidine deaminases (Alexandrov et al., 2013). Breast cancer samples without any mutations recorded as either signatures 2 or 13 were classified as being without AID/APOBEC mutation signatures.

### Methylation data

Methylation data from normal sigmoid colon tissue was downloaded from the Roadmap Epigenomics Atlas (Roadmap Epigenomics Consortium et al., 2015) (Gene Expression Omnibus [GEO]: GSM983645). These data were from WGBS, and were obtained as a wig file, converted to BED format using “convert2bed”. Methylation values and chromosome coordinates for individual nucleotides in each CpG were merged, taking the methylation value for the cytosine of each CpG dinucleotide. This value was then used for all methylation calculations (when matched with colon cancer sample mutations) throughout this study.

Additional methylation datasets were obtained from the Roadmap Epigenomics Atlas (Roadmap Epigenomics Consortium et al., 2015) and analysed similarly. These datasets were matched to various cancer types and subtypes as listed in **Supp Table 1**, together with their GEO accession numbers.

Where data are binned across methylation values, bins spanning 0.1 methylation were used for all methylation values between 0 and 1. For CpGs with methylation values equalling exactly 0 or 1, data were allocated to bins representing either 0 or 1 methylation only, respectively.

Regression models and correlations which incorporated methylation data (that is, **Fig 1b**, **Fig 5**, **Fig 6**, **Supp Fig 2a**, **Supp Fig 5** and **Supp Fig 6**) used only methylation values from CpG dinucleotides located on autosomes.

### Promoter profiles

H3K4me3 chromatin immunoprecipitation sequencing (ChIP-seq) data from normal sigmoid colon tissue were obtained from the Roadmap Epigenomics Atlas (Roadmap Epigenomics Consortium et al., 2015) (GEO: GSM956024), and converted to bigwig using “wigToBigWig”. Normal sigmoid colon tissue WGBS methylation data were obtained as described above, and converted to bigwig similarly. DNase I hypersensitivity (DHS) data were obtained for HCT116 cells from the ENCODE project consortium (The Encode Project Consortium, 2012) and downloaded as a bigwig file through UCSC (GEO: GSM736600).

Transcription start sites (TSSs) for each gene were obtained from the UCSC table browser. Mutation profiles were generated by counting mutations at each base within ± 4 kb of a TSS, with counts normalised to mutations per Mb. Methylation, H3K4me3 and DHS profile data were created by use of the “computeMatrix” (reference-point) and “plotProfile” tools available through the deepTools package (Ramírez et al., 2014). All profiles were orientated so that the gene body runs from 5’ to 3’, downstream from the TSS.

### Replication timing analyses

Genome-wide replication timing datasets were downloaded from UCSC Genome Browser (also available through GEO as GSE34399). GM12878 was the only lymphoblastoid cell-line used, to avoid biasing the sample through inclusion of multiple lymphoblastoid cell-lines, as previously described (Supek and Lehner, 2015). The remaining datasets contained replication timing values for 11 cell-types. The genome was divided into megabase windows using BEDtools (Quinlan and Hall, 2010), with replication timing averaged across cell-types within these windows. Thus data points in **Supp Fig 2a** show average replication timing and methylation in megabase bins. To ensure robust measurements with enough data points, sites were excluded from some analyses if they had replication timing listed as <20 or >80. (Replication timing values in the raw data range from 0 to 100. For presentation in figures, these values have been inverted, such that lower values indicate earlier replication). chrY was excluded from replication timing analyses, as values for this chromosome were not present in the original raw data.

MSI-signature mutations were determined using the “MSI-enriched signatures” from Supek & Lehner (2015) “Extended Data Figure 4i”. Therefore, non-C[>T]pG MSI signature mutations include all of the following, in addition to their complementary trinucleotides and alternate nucleotide: C[C>A]A, C[C>A]T, C[C>A]C, C[C>A]G, A[C>T]A, A[C>T]T, A[C>T]C, G[C>T]A, G[C>T]T, G[C>T]C, T[A>G]A, T[A>G]C, T[A>G]T, T[A>G]G, C[A>G]A, C[A>G]C, C[A>G]T, C[A>G]G, A[A>G]A, A[A>G]T, A[A>G]C, A[A>G]G, G[A>G]A, G[A>G]C, G[A>G]T, and G[A>G]G. C>T CpG mutations included A[C>T]G, C[C>T]G, G[C>T]G and T[C>T]G, together with their complementary trinucleotides (for G>A mutations only). Full MSI-signature mutations included all trinucleotide/mutation options listed above. This signature corresponds closely with signature 6 from Alexandrov et al. (2013), which is associated with microsatellite instability.

Excision repair sequencing (XR-seq) data for skin fibroblast cell line NHF1 (Hu et al., 2015) were obtained in Sequence Read Archive (SRA) format (GEO: GSM1659156), and processed as previously described (Poulos et al., 2016).

### Strand specificity and origins of replication

*TOP1* and *LMNB2* oriC sites were selected for use since they were well-defined oriC per Shinbrot et al. (2014). The region used in analyses as that surrounding the oriC for *TOP1* was chr20:39,300,000-39,900,000, with the oriC isolated upstream of the *TOP1* TSS as shown in **Fig 3d** (upper panel). The region used in analyses as that surrounding the oriC for *LMNB2* was chr19:2,000,000-2,700,000, with the oriC as given in Shinbrot et al. (2014) and shown in **
Fig 3d** (lower panel).

### Statistical analyses

Regression models and other statistical analyses were performed in R. For each cancer type or subtype, the binary logistic regression model used incorporated methylation (with a quadratic term), replication timing and an interaction between methylation and replication timing, as shown below:

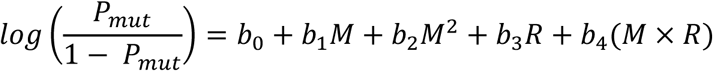

*Where, P_mut_*: =*probability of mutation*
*M*: =*methylation*
*R*: =*replication timing*
*b_0_, b_1_, b_2_, b_3_ and b_4_*: *represent constants estimated from logistic regression*.

This model was selected for use as it significantly improved upon nested binary logistic regression models with fewer terms (data not shown). A significant improvement was determined by use of both a Likelihood Ratio Test (LRT; “lrtest” function from the “lmtest” package (Zeileis and Hothorn, 2002); model selected if LRT showed significant improvement by *P* < 0.05 at all steps between nested models) and the Akaike Information Criterion (AIC; model with smallest AIC was selected).

Regression models were constructed using data for autosomes only. Mutations were considered a binary outcome, with each CpG designated as either never mutated in any sample, or mutated in at least one sample, within a given cancer type or subtype. The area under the curve (AUC) was calculated using the ROCR package (Sing et al., 2005).

Equations predicted by the regression models, together with the predicted vertex and AUC from relevant nested models are recorded in **Table 1** and **Supp Table 2**. To plot the actual and predicted values from the regression model (as in **Fig 5**, **Fig 6** and **Supp Fig 5**), data was binned either by methylation (bin size of 0.1, ranging from 0 to 1; for CpGs with methylation values equalling exactly 0 or 1, mutations were allocated to bins representing 0 or 1 methylation only, respectively) or by replication timing (bin size of 10, ranging from 20 to 80).

Regarding **Fig 5** and **Supp Fig 5**, where mutation probability or log odds of mutation probability was plotted against methylation (leftmost two graphs, respectively), an average was used for replication timing within each bin. To separate the influence of each factor, where the predicted function was plotted against methylation using average replication timing (middle graph) or against replication timing using average methylation (rightmost graph), the overall genome-wide average for replication timing or methylation (respectively) was used in the equation for all bins. Where log odds of mutation probability was plotted against replication timing (remaining graph), an average was used for methylation within each bin.

In **Fig 1b**, significance was determined by Pearson’s correlation on binned data, with slopes of MSI and *POLE*-mutant colorectal cancer compared to MSS colorectal cancer data via a Poisson regression, with MSS as the factor level reference. Statistical analyses shown in **Fig 2a**, **b** and **Supp Fig 2d** were performed using Pearson’s correlation on binned data. Statistical analyses displayed in **Supp Fig 2a**, **c**, **
Fig 3a** and **Supp Fig 6a** were performed using Pearson’s correlation on the data points shown. Fisher’s exact test was used to determine the levels of significance given in **Fig 3c
** and **Fig 4a**. **
Fig 3b** shows significance by paired t-test between samples and **Fig 4b** displays significance by one-sample t-test. All other determinations of significance were made by unpaired t-test. In all instances, significance was determined using a threshold of *P* < 0.05.

## Acknowledgements

The authors thank TCGA and other groups who have made data publically available. This work was funded by the National Health and Medical Research Council (NHMRC, Australia) (APP1119932), Cancer Institute NSW (13/DATA/1-02) and the Cure Cancer Foundation Australia with the assistance of Cancer Australia through the Priority-driven Collaborative Cancer Research Scheme (APP1057921) to J.W.H.W. R.C.P is supported by an Australian Postgraduate Award.

---

**Supplementary Figure 1.**
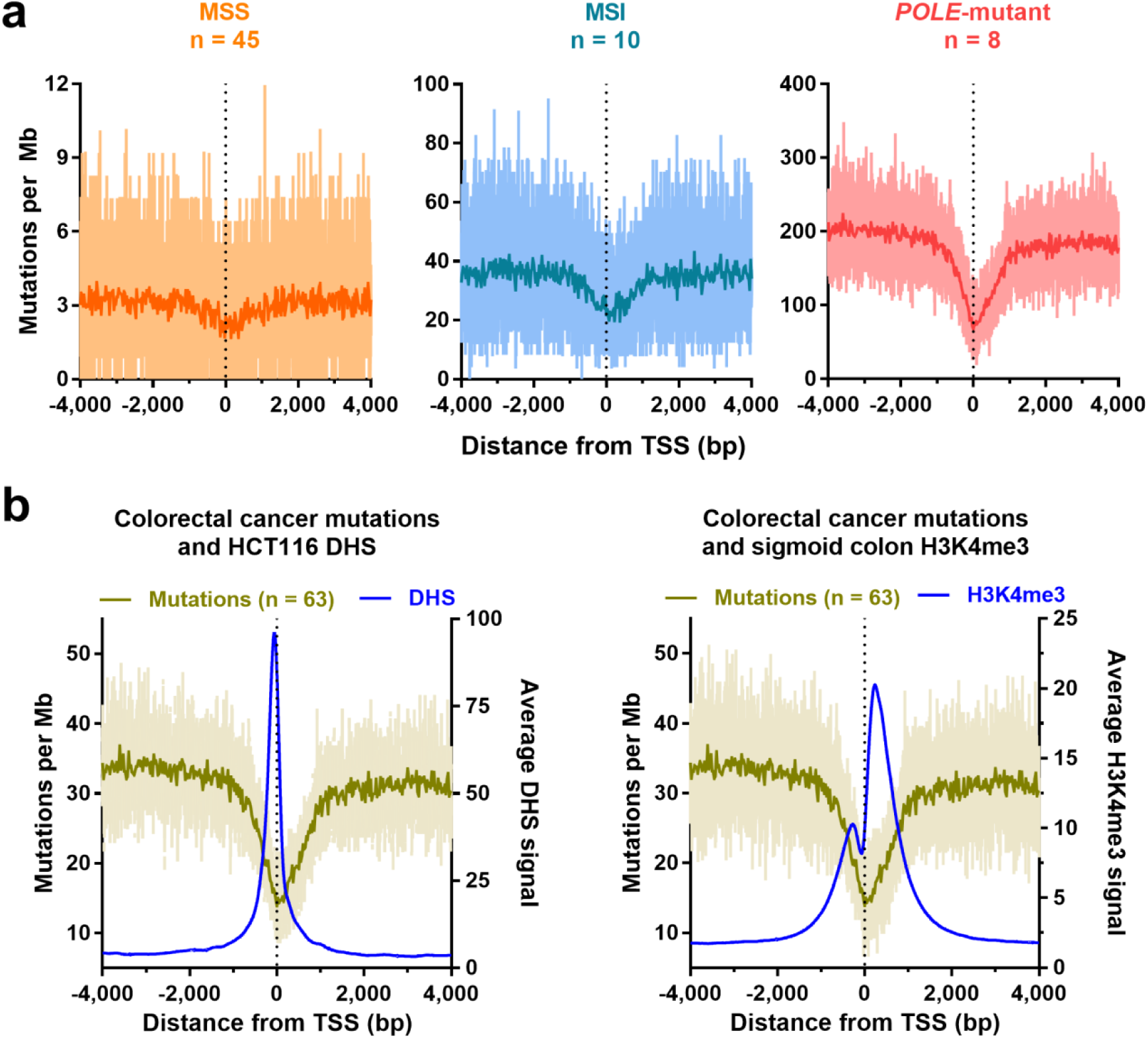
**Mutation, DNase I hypersensitivity (DHS) and H3K4me3 signal around transcription start sites (TSSs) in colorectal cancer. (a)** Mutation profiles around the TSS for microsatellite stable (MSS) colorectal cancer (left panel), those with microsatellite instability (middle panel) and with *Polymerase epsilon* exonuclease domain mutation (*POLE*-mutant; right panel). For mutations, nucleotide-resolution data is shown (light colour) along with data in 25 bp bins (dark colour). (b) Colorectal cancer mutation profile along with average DHS signal from HCT116 colorectal cancer cell-line (left panel) or colon tissue H3K4me3 signal (right panel) around the TSS.

**Supplementary Figure 2.**
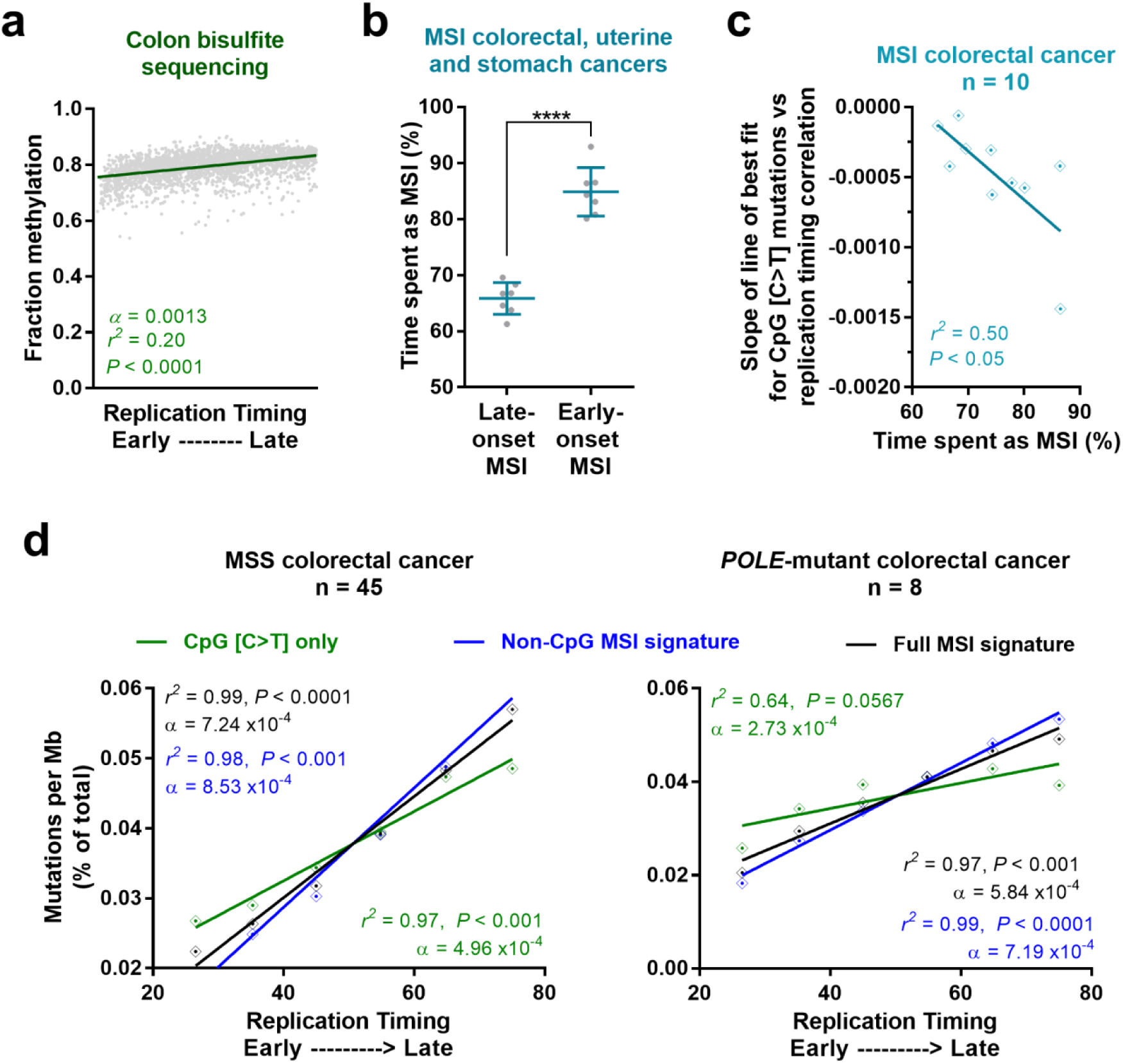
**Replication timing, methylation and MSI-associated mutations in colorectal cancer subtypes. (a)** Association between normal sigmoid colon methylation from whole-genome bisulfite sequencing and replication timing across autosomes. Grey dots show data binned to megabase-scales. **(b)** Time spent with microsatellite instability (MSI) (%) for late– and early-onset MSI uterine, stomach and colorectal cancer samples. Significance is by unpaired t-test, where **** P < 0.0001. **(c)** Association between the slope of the line of best fit from CpG C>T mutations versus replication timing-association, and time spent as MSI for colorectal cancer samples. **(d)** Association between CpG C>T mutations, non-C[>T]pG MSI signature and full MSI signature (including CpG C>T) mutations with replication timing for microsatellite stable (MSS; left panel) colorectal cancers and colorectal cancers with *Polymerase epsilon* exonuclease domain mutation (*POLE*-mutant; right panel). Data points show binned data (bins of 10 replication timing). Unless otherwise stated, *r*^*2*^ and significance are by Pearson’s correlation and α denotes the slope of the line.

**Supplementary Figure 3.**
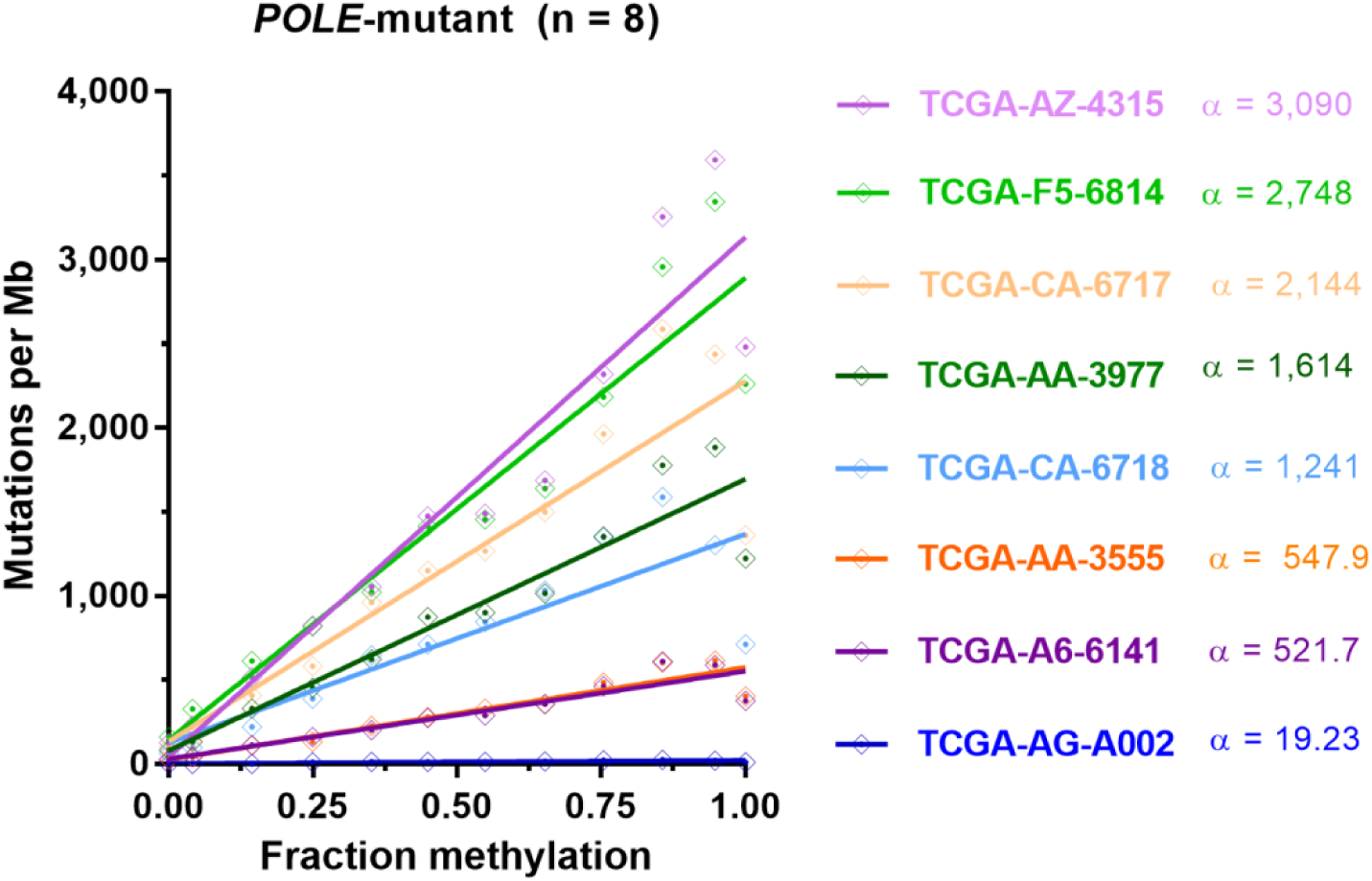
**Association between mutations and methylation in individual *POLE*-mutant colorectal cancers.** Line of best fit from binned data of mutation-methylation associations in *Polymerase epsilon* exonuclease domain mutant (*POLE*-mutant) colorectal cancers. α denotes the slope of the line of best fit, with data binned by 0.1 methylation.

**Supplementary Figure 4.**
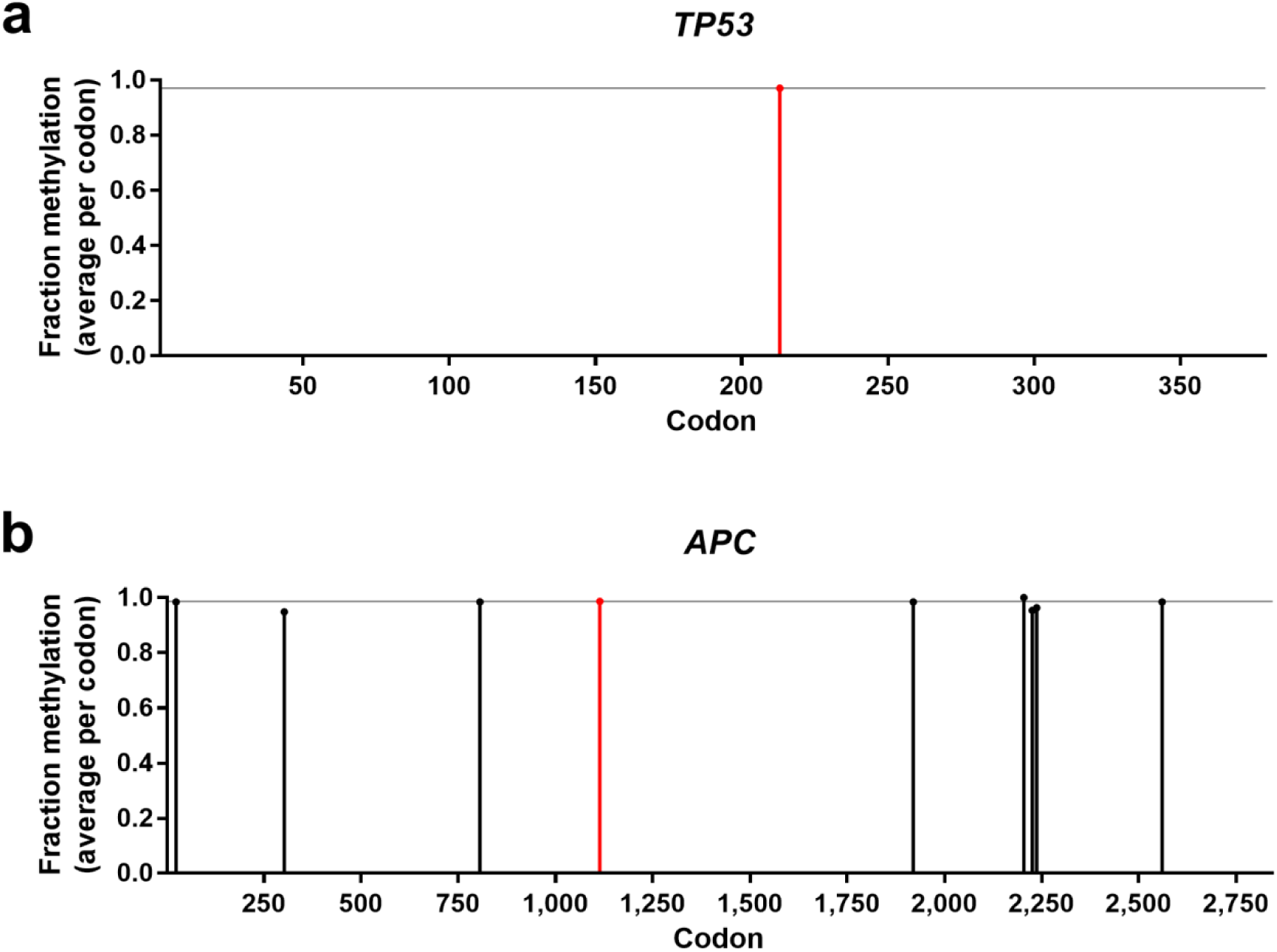
**Methylation status for all possible sites of truncating T[C>T]G mutations within *TP53* and *APC* genes in normal colon tissue.** Methylation status in normal colon tissue of each TCG trinucleotide which, via a C>T mutation, would result in a protein truncation within a coding exon of *TP53* (top) and *APC* (bottom). The R213 (*TP53*) and R1114 (*APC*) codons are indicated in red, with a horizontal line marking the methylation level at these codons.

**Supplementary Figure 5.**
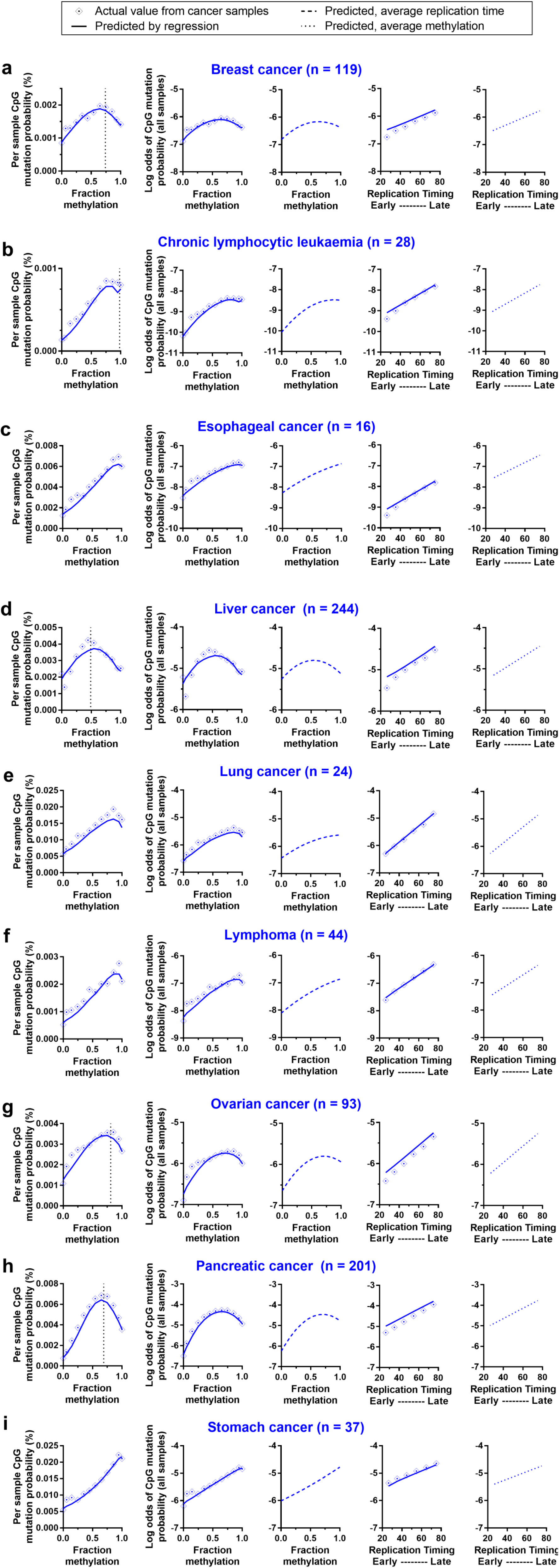
**Actual and predicted mutation rates, according to methylation and replication timing, across cancer types.** Graphs depict actual and predicted (by regression model; see Methods) mutation probability and log odds of mutation probability by methylation or replication timing, for (a) breast cancer, (b) chronic lymphocytic leukaemia, (c) esophageal cancer, (d) liver cancer, (e) lung cancer, (f) lymphoma, (g) ovarian cancer, (h) pancreatic cancer and (i) stomach cancer. Graphs from left to right are: mutation probability by fraction methylation (actual and predicted), log odds of mutation probability by fraction methylation (actual and predicted), log odds of mutation probability by fraction methylation (predicted, using overall average replication timing in all bins), log odds of mutation probability by replication timing (actual and predicted) and log odds of mutation probability by replication timing (predicted, using overall average methylation in all bins). Binned data is shown (bins of 0.1 for methylation or 10 for replication timing), with any vertex between 0 and 1 fraction methylation indicated by a dotted line. See Supp Table 2 for regression output, predicted vertex and area under curve values.

**Supplementary Figure 6.**
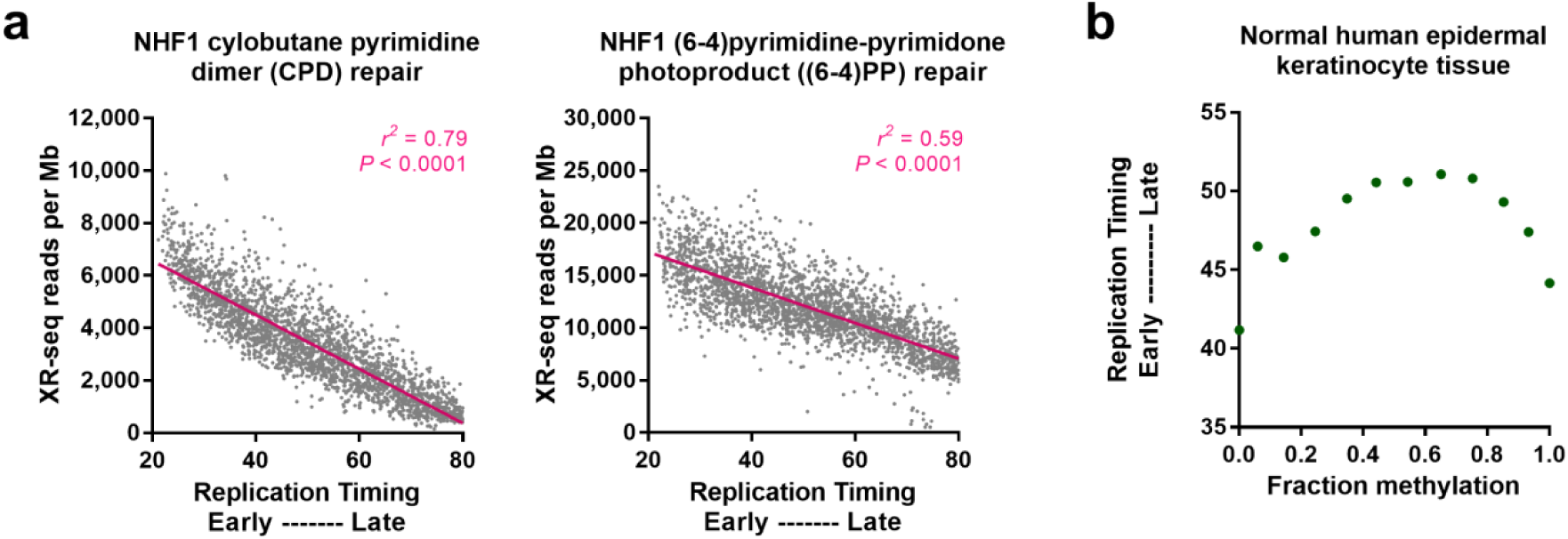
**Associations with replication timing of nucleotide excision repair and methylation in skin cells. (a)** Association between cyclobutane pyrimidine dimer (CPD; left) or (6-4)pyrimidine-primidone photoproduct ((6-4)PP; right) excision sequencing (XR-seq) repair reads with replication timing. *r*^*2*^ and significance is by Pearson’s correlation, with grey dots showing data binned to megabase scales. (b) Association between replication timing and normal human epidermal keratinocyte (NHEK) cell methylation. Data is shown in bins of 0.1 methylation.

## Supplementary Tables

**Supplementary Table 1.** Methylation datasets matched with each cancer type and subtype used in regression analyses.

| Cancer Type | Samples* | WGBS tissue type<sup>#</sup> | GEO<sup>+</sup> accession |
| --- | --- | --- | --- |
| Breast cancer | 119 | Breast luminal epithelial cells | GSM1127125 |
| Breast cancer:<br>without AID/APOBEC<br>signatures | 88 | Breast luminal<br>epithelial cells | GSM1127125 |
| Breast cancer:<br>with AID/APOBEC signatures | 14 | Breast luminal<br>epithelial cells | GSM1127125 |
| Chronic lymphocytic leukaemia | 28 | CD34, mobilized<br>primary cells | GSM916052 |
| Esophageal cancer | 16 | Esophagus, adult | GSM983649 |
| Liver cancer | 244 | Liver, adult | GSM916049 |
| Lung cancer | 24 | Lung, adult | GSM983647 |
| Lymphoma | 44 | Thymus | GSM1010979 |
| Melanoma | 36 | Foreskin keratinocyte<br>primary cells | GSM1127056 |
| Ovarian cancer | 93 | Ovary, adult | GSM1010980 |
| Pancreatic cancer | 201 | Pancreas | GSM983651 |
| Squamous cell carcinoma (all<br>samples are XPC<sup>-/-</sup>) | 5 | Foreskin keratinocyte<br>primary cells | GSM1127056 |
| Stomach cancer | 37 | Gastric, adult | GSM1010984 |

\* All samples are from The Cancer Genome Atlas (TCGA), with the exception of the XPC<sup>-/-</sup> squamous cell carcinomas which were published by Zheng et al. (2014).
<sup>#</sup> WGBS = whole genome bisulfite sequencing. All tissue-types used were from normal cells.
<sup>+</sup> GEO = Gene Expression Omnibus

**Supplementary Table 2.** Regression equation from multivariable models predicting mutation probability across cancer types and subtypes, together with vertex and area under curve (AUC) predictions.

| **Cancer type/subtype** | **Regression model equation\*** | **Vertex<sup>^</sup>**<br>**(fraction methylation)** | **AUC<sup>@</sup> for regression model A<sup>A</sup>** | **AUC<sup>+</sup> for regression model B<sup>B</sup>** | **AUC<sup>+</sup> for regression model C<sup>C</sup>** | **Increase to AUC via replication<sup>#</sup>** | **Increase to AUC via methylation<sup>+</sup>** |
| --- | --- | --- | --- | --- | --- | --- | --- |
| Breast cancer: with AID/APOBEC sig. | $y = -7.7013 + 1.1657M + -1.1587M^2 + -0.0019*R + -0.0098(M \times R)$ | 0.50 | 0.562 | 0.544 | 0.576 | 2.6% | 6.0% |
| Breast cancer without AID/APOBEC sig. | $y = -6.7419 + 2.8769M + -1.9303M^2 + -0.0142*R + -0.0035(M \times R)$ | 0.75 | 0.560 | 0.594 | 0.610 | 8.8% | 2.6% |
| Breast cancer | $y = -6.2782 + 2.3895M + -1.6076M^2 + -0.01*R + -0.0069(M \times R)$ | 0.74 | 0.550 | 0.584 | 0.594 | 8.0% | 1.7% |
| Chronic lymphocytic leukaemia<sup>=</sup> | $y = -8.9527 + 3.9184M + -2.0027M^2 + -0.0212*R + -0.0072(M \times R)$ | 0.98 | 0.557 | 0.649 | 0.666 | 19.7% | 2.7% |
| Esophageal cancer<sup>=</sup> | $y = -7.1719 + 2.1664M + -0.5986M^2 + -0.0211*R + -0.0031(M \times R)$ | 1.81 | 0.561 | 0.622 | 0.644 | 14.8% | 3.5% |
| Liver cancer | $y = -4.3453 + 1.5195M + -1.5371M^2 + -0.0172*R + 0.0029(M \times R)$ | 0.49 | 0.555 | 0.585 | 0.593 | 6.8% | 1.3% |
| Lung cancer | $y = -5.2744 + 1.9996M + -0.7005M^2 + -0.0221*R + -0.0087(M \times R)$ | 1.43 | 0.552 | 0.648 | 0.657 | 19.0% | 1.4% |
| Lymphoma | $y = -7.1358 + 2.09M + -0.565M^2 + -0.0185*R + -0.0055(M \times R)$ | 1.85 | 0.564 | 0.624 | 0.639 | 13.4% | 2.5% |
| Ovarian cancer | $y = -5.8518 + 2.7244M + -1.6735M^2 + -0.015*R + -0.0068(M \times R)$ | 0.81 | 0.552 | 0.608 | 0.620 | 12.3% | 2.0% |
| Pancreatic cancer | $y = -4.8767 + 4.9924M + -3.6038M^2 + -0.026*R + 0.0015(M \times R)$ | 0.69 | 0.592 | 0.634 | 0.658 | 11.0% | 3.7% |
| Stomach cancer | $y = -5.2204 + 0.9192M + 0.2171M^2 + -0.0151*R + 0.0018(M \times R)$ | - 2.12 | 0.573 | 0.571 | 0.606 | 5.8% | 6.3% |

\* See **Methods** for canonical regression model formula, where $y$ = log odds of mutation probability, $M$ = methylation and $R$ = replication timing
<sup>^</sup> Vertex (unit: fraction methylation) predicted by regression model, calculated as $-b_1/(2 \times b_2)$ .
@ AUC = area under curve
<sup>A</sup> Regression model equation: $y = b_0 + b_1 M + b_2 M^2$ where $y$ = log odds of mutation probability, $M$ = methylation and $R$ = replication timing
<sup>B</sup> Regression model equation: $y = b_0 + b_1 R$ where $y$ = log odds of mutation probability, $M$ = methylation and $R$ = replication timing
<sup>C</sup> Regression model equation: $y = b_0 + b_1 M + b_2 M^2 + b_3 R$ , where $y$ = log odds of mutation probability, $M$ = methylation and $R$ = replication timing
<sup>#</sup> Calculated using values of AUC from models: $(C-A)/A$
<sup>+</sup> Calculated using values of AUC from models: $(C-B)/B$
<sup>=</sup> For both chronic lymphocytic leukaemia (CLL) and esophageal cancers, the LRT did not indicate that there was a significant improvement between the model as given above but excluding the interaction term ( $M \times R$ ), and the full model as stated above, including the interaction term ( $P = 0.1303$ and $P = 0.1055$ , respectively). However, as the AIC was still smallest for the full model which included the interaction term, and for consistency of model usage across cancer types, the full model was used for the analyses of CLL and esophageal cancer.

